# Code Multiplexed Nanocapacitor Arrays for Scalable Neural Recordings

**DOI:** 10.1101/2024.02.22.581578

**Authors:** Sean Weaver, Yiyang Chen, Aline Renz, Woojun Choi, Yashwanth Vyza, Tilman Schlotter, Katarina Vulić, Donghwan Kim, Gabriele Atzeni, Dmitry Momotenko, Nako Nakatsuka, Taekwang Jang, János Vörös

**Affiliations:** Laboratory for Biosensors and Bioelectronics, ETH Zürich; Zürich, 8092, Switzerland; Energy Efficient Circuits and IoT Systems Group, ETH Zürich; Zürich, 8092, Switzerland; Mixed-Signal Integrated Circuits and Systems Laboratory, KHU; Yongin-si, 17104, Korea

## Abstract

Large scale neural recordings are redefining our understanding of the brain. However, simultaneously recording potentials from thousands of microelectrodes remains challenging. We overcome this limitation by measuring activity-induced changes in mutual capacitance. Code division multiplexing enabled simultaneous recordings of neural activity from 1,024 nanocapacitor electrodes at a density of 10k electrodes/mm.^2^ Features of neural activity *i*.*e*., action potentials, bursts, and local field potentials were measured in recordings, benchmarking our device against the state-of-the-art.

**Summary:** Capacitive sensing of neural activity improves the scalability, fabrication, and miniaturization of microelectrode arrays.

## Main Text

Measuring neural activity across the brain with high spatial and temporal resolution is a prerequisite to answering questions in neuroscience. The last 30 years have seen advancements in optical tools, *e*.*g*., high-speed recordings of neural ensembles (*1*), and microelectrode arrays, both *in vivo* (*2, 3*), and *in vitro* (*4, 5*). Despite reports of advances in optical recordings, (*6, 7*) fundamental principles underlying electrophysiology sensors have not expanded since the introduction of field-effect transistor (FET)-assisted recordings in 1981 (*8*). The extent to which FET or passive microelectrode arrays (MEAs) will meet experimental demands remains unclear. Brain-wide recordings with single neuron resolution require electrodes suitable for dense implantation, power consumption of ∼0.2 μW/electrode, and simultaneous sampling of ≳100k channels for small mammals (*9, 10*). Research on electrode material (*11*– *14*), geometry (*15*–*18*), and recording electronics (*19, 20*) has improved performance, but conventional MEAs measuring potential scale unfavorably as electrodes are miniaturized since this leads to higher impedance, increased noise, power consumption, and/or circuit size. Capacitive sensing can compensate impedance increases associated with miniaturization. Similar to FET arrays (*21, 22*), capacitive sensors can be used with frequency or code division multiplexing (FDM or CDM) for scalable parallel readout. To this end, we present the theoretical basis for and practical demonstration of mutual capacitance measurements of neural activity. Our application specific integrated circuit (ASIC) with 32 drivers and 32 amplifiers sampled an array of 1,024 nanogap capacitors using CDM at a density of 10k electrodes/mm^2^ detecting both action potentials (APs) and local field potentials (LFPs).

## Results

Sensor design requires a conceptually simple, but physically accurate, description of the transducer. Direct (Fig. 1Ai) or FET-assisted (Fig. 1Aii) surface potential measurements model a neuron as a voltage source capacitively coupled to the electrode: however, inclusion of circuit approximations for the Nernst-Plank-Poisson (NPP) equations are favored in practice (*23*). Mutual capacitive measurements of neural activity would treat cell-driven changes in ion concentration as a variable capacitance connected to ground situated between electrodes of opposite potential (Fig. 1B). To conceptually validate this simple model and investigate how cleft distance impacts signal amplitude we performed finite element simulations (fig. S1) coupling a Hodgkin-Huxley (*24*) neuron model to the Nernst-Planck equations for electrodiffusion (*25*) The model includes electrodynamic, as opposed to electrostatic (*26*), simulations of the intra/extra-cellular space. Membrane conductance, flux, and ion relevant constants were based on the electrodiffusive Pinsky-Rinzel model (*27*). Potentials were evaluated at the inner/outer membrane (Fig. 1Ci/Cii), at the electrode (Fig. 1C*iii*), and compared to the Goldman-Hodgkin-Katz (GHK) reversal potential (Fig. 1Civ). Concentration of major ionic species and potential at the electrode during 3 APs varied with cleft height (Fig. 1D). Concentration differences (*e*.*g*., 2.6 mM *vs*. 0.7 mM Na^+^) at the electrode for small changes in distance (*e*.*g*., 10 nm *vs*. 40 nm) highlight the importance of the cell-electrode interface. Potential and charge density vary minimally across the distance between the membrane and electrode (fig. S2). Greater impact on sensitivity is due to total cleft volume (fig. S3). During an AP, capacitance at the electrode surface varies (Fig. 1E, fig. S4) due to changes in ion concentration within the sensing volume. At a potential of 20 μV, the change in capacitance of the first element above the electrode is either -2 nF/μm^3^ for the rising or -1.75 nF/μm^3^ for the falling edge of an AP (fig. S5). An AC signal between electrodes at low potentials and high frequencies should have minimal impact on the double layer (*28*) (fig. S7, S8). Validation experiments are shown in fig. S9.

**Figure 1:**
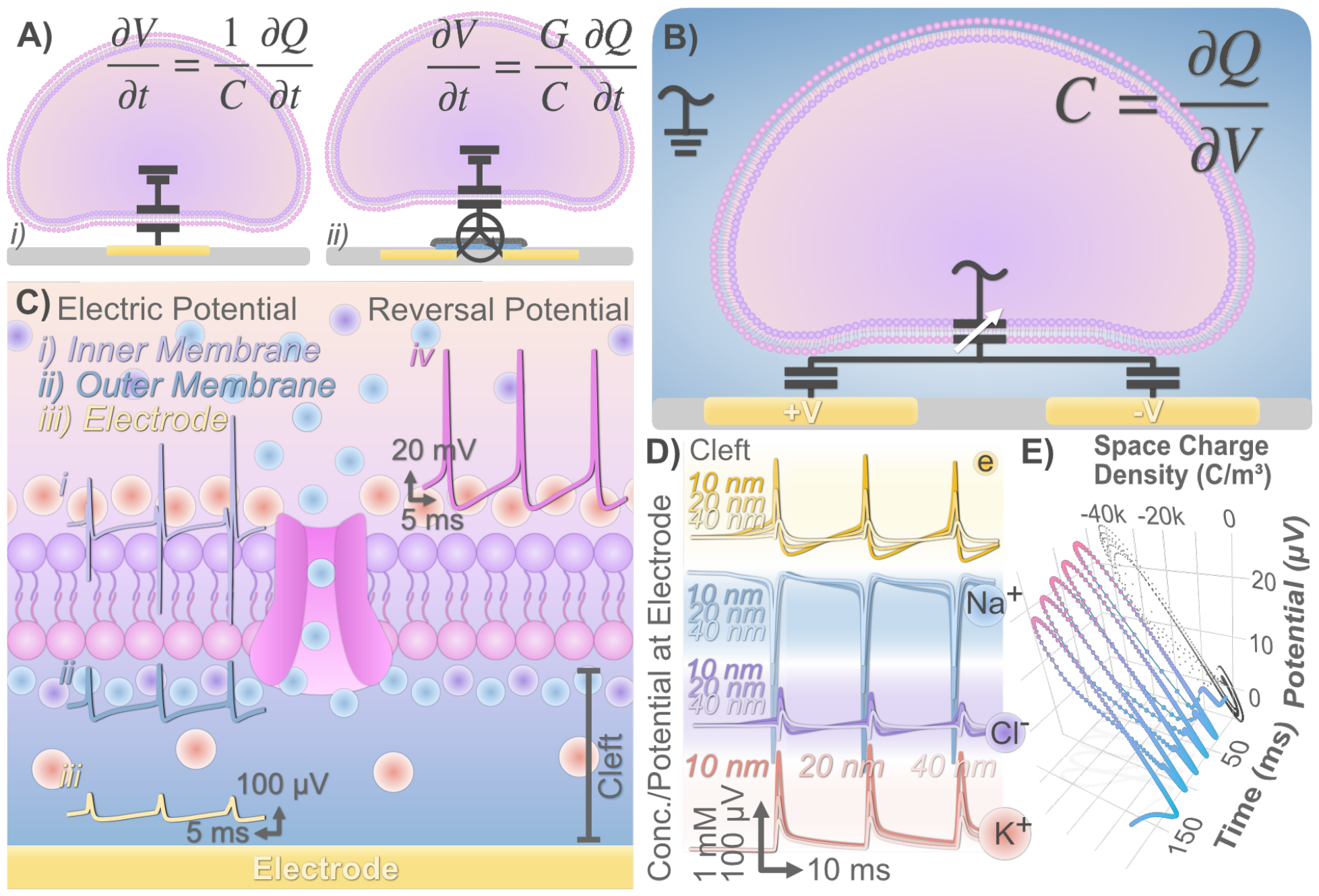
Principle and simulation of mutual capacitive sensing of neural action potentials. (**A**) Circuit model of two methods dominating extracellular recordings i) sensing voltage, V induced by changes in ion concentration, *i*.*e*., charge Q, or ii) FET measurements amplifying potential changes by a gain, G. The neuron acts as a potential source by transiently violating electroneutrality. (B) Circuit model for mutual capacitive measurements of neural activity. Neural activity adds/removes charge from the effective dielectric between the electrodes acting as a variable capacitor connected to ground. An alternate circuit model is shown in fig. S6. (C) Electric potential derived from local current densities of a Hodgkin-Huxley neuron at the i) inner membrane wall, ii) the outer membrane wall, iii) and the electrode surface as compared to the GHK reversal potential (Cleft = 20nm). (D) Changes in [K^+^], [Cl^-^], [Na^+^] and potential for Cleft = 10, 20, 40 nm across 3 APs at the electrode. (E) Relation between space charge density (i.e. charge concentration) and potential across 5 APs. Non-unique values for charge at a given potential indicate a change in electrode capacitance through the course of an action potential (Cleft = 40 nm). Projection of potential *vs*. charge through time is in black.

Mutual capacitance sensors are suited to multiplexing schemes allowing concurrent readout of all elements. In CDM, orthogonal *i*.*e*., separable, digital signals are sent to one electrode of each capacitor sharing a common second (Fig. 2A). We used Walsh coding to generate orthogonal signals. The receiving plate (Rx) potential (resp. current for transimpedance) is the superposition of capacitively coupled signals from each transmission electrode (Tx). Demodulating each Rx by multiplication with the orthogonal codes reconstructs electrode capacitances. This method of multiplexing substantially reduces the number of amplifiers, interconnects, and routing, while avoiding the bandwidth constraints of high channel count FDM. Nanogap capacitor (*29, 30*) arrays (Fig. 2B) can be made using standard microfabrication techniques on various substrates, in contrast to CMOS, with similar electrode count and density. Our MEAs had 32 × 32 Au tracks with a pitch of 10 μm and a width of 3 μm separated by 200 nm SiO_2_ (Fig. 2C, fig. S10). A linear response to changes in osmolarity of 0.2 pF/mOsm was found between 240 –300 mOsm (Fig. 2D), the expected range for neural cultures (neurobasal media (NB): 260±10 mOsm, ∼300mOsm for human cerebrospinal fluid). During an AP, our simulation results indicate osmolarity changes on the order of 0.5 – 5 mOsm depending on neuron-electrode coupling. The input referred noise of the ASIC has an equivalent capacitive noise of 39 – 104 fF (root-mean-square). The lower end of these changes is at ASIC noise floor edge (*31*).

**Figure 2:**
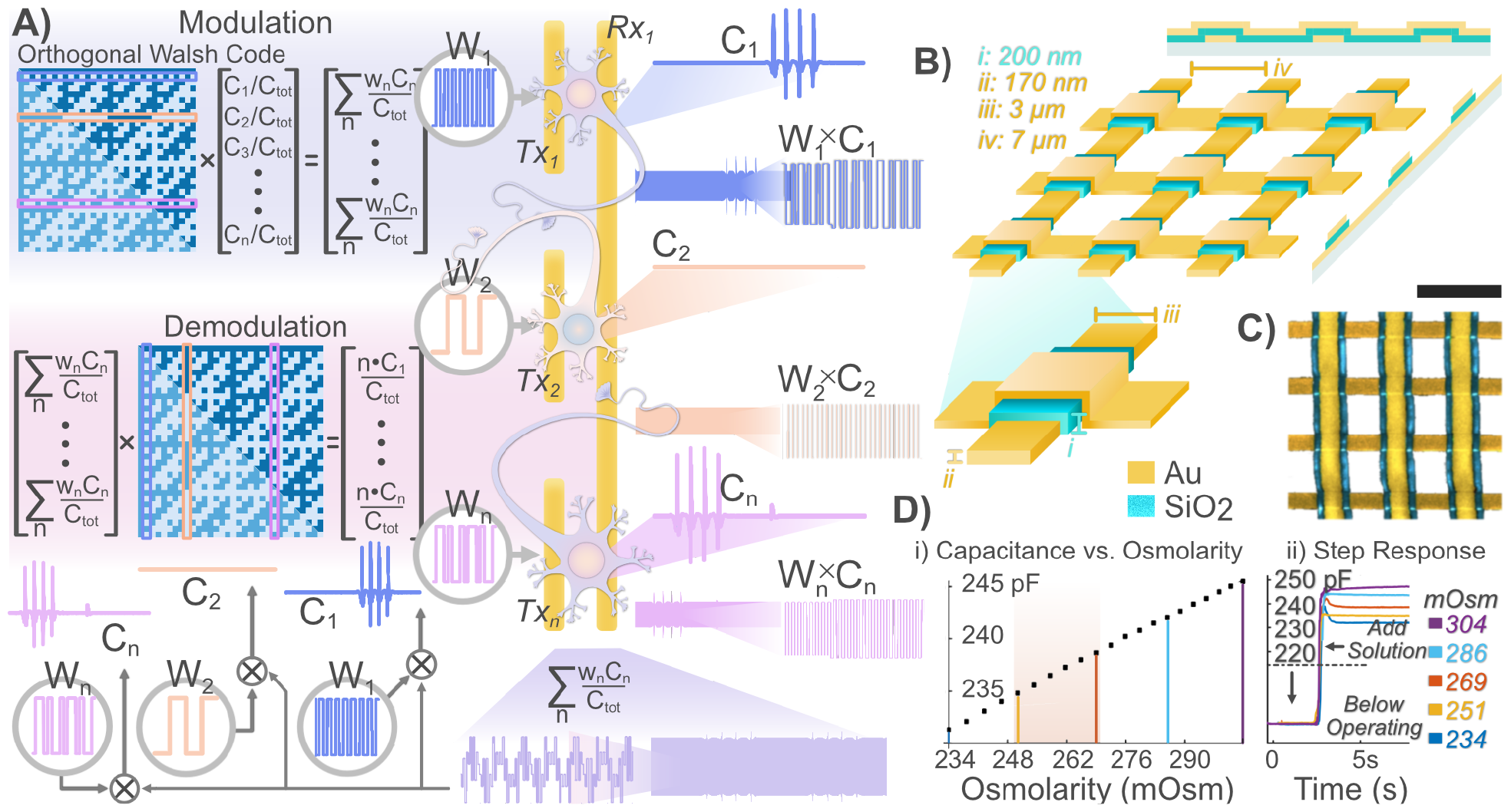
Code division multiplexed readout of nanocapacitor arrays. (**A**) 32-bit binary orthogonal codes are generated using the Walsh algorithm. Each bit is translated to a 1.56 μs long potential within a 50 μs long code (W_n_) applied to condenser plates (Tx) comprising half of 32 independent capacitors. Capacitance changes induced by neurons (C_n_) on the array are convolved with the codes producing a combined modulation signal on a common receiving condenser plate (Rx). The modulated signal is a linear combination of 32 capacitances. C_tot_ is the total capacitance of each Rx. Multiplying the modulated signal with each original code reconstructs the capacitance for each Rx capacitor. (B) Nanocapacitor arrays are fabricated by depositing two gold layers on glass separated by a 200 nm thick layer of SiO_2_ (i). Gold tracks are 170 nm thick (ii) and 3 μm wide (iii). Edge-to-edge spacing between tracks is 7 μm (iv). (C) A false colored SEM image of the fabricated array (scale = 10 μm). (D) Changes in capacitance due to variation in osmolarity for i) serial dilution for a dose response curve (NB osmolarity range in orange) and ii) injection for the step response of the sensor starting from a baseline in MilliQ water.

Nanogap capacitors have not previously been used for neural recordings, so we studied the neuron-electrode interface using focused ion beams (FIB) to mill nanocapacitor cross sections subsequently imaged by a scanning electron microscope (SEM). Neurites from day *in vitro* (DIV) 21 rat hippocampal culture growing across a 1 × 32 nanocapacitor array showed no avoidance of the electrode surface (Fig. 3A, SEM of 32×32 arrays shown in fig. S11). Analysis of 30 cross sections from 4 samples showed an average vertical distance from the electrode of <10 nm for 8 repeats while 12 had an average distance >100 nm (Fig. 3B). Neurons often encapsulated the nanocapacitor (Fig. 3C, fig. S12) or formed cavities across the surface (Fig. 4D). Engulfment of electrodes has been shown in other work to increase signal strength (*32*) due to larger local ion concentrations during APs given decreased diffusion to bulk within the sensing region.

**Figure 3:**
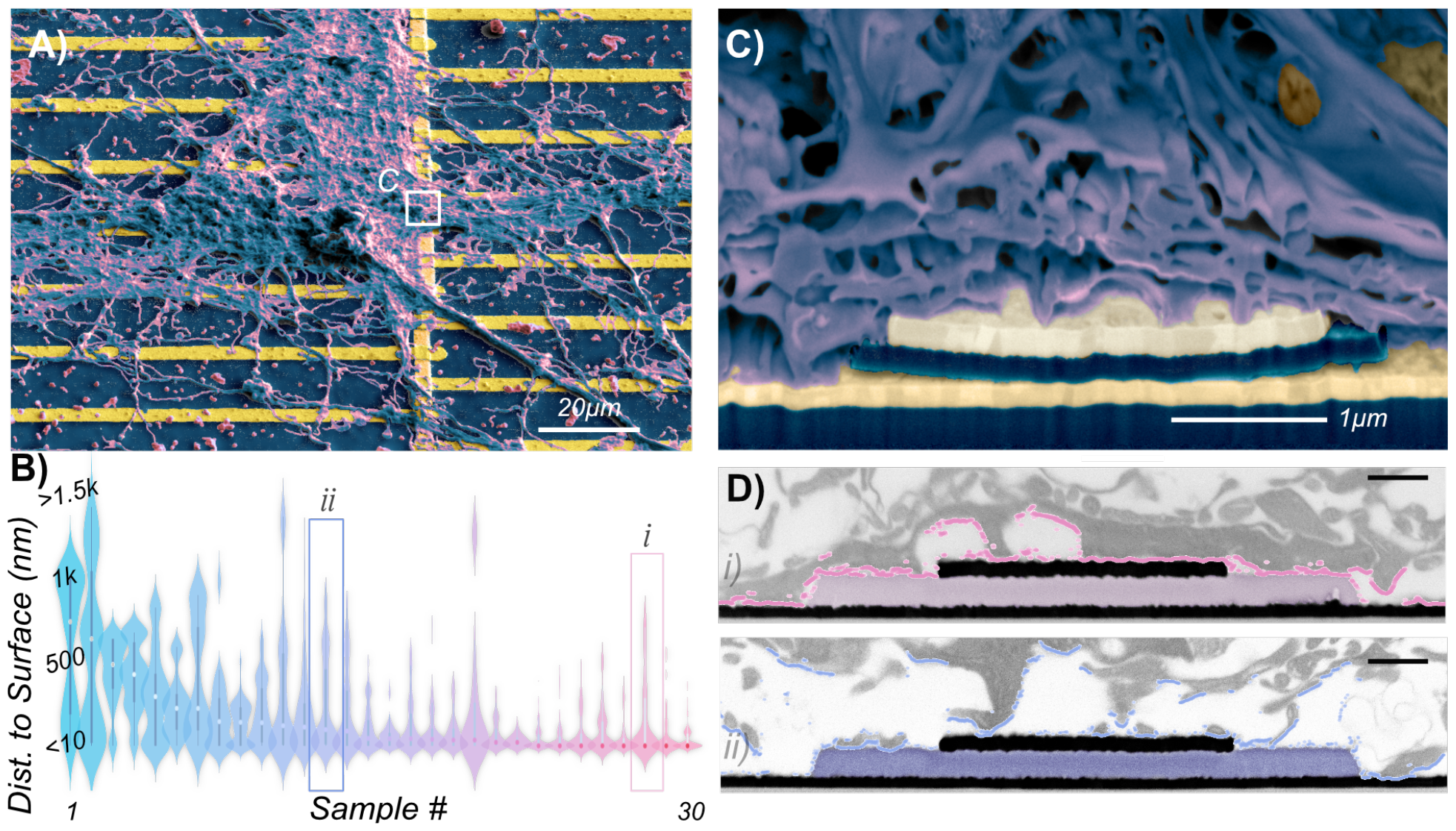
Neuron-nanocapacitor interface. (**A**) SEM of ultra-thin plasticized neurons on top of a 1×32 channel nanocapacitor array (B) Estimates of the vertical distance between the nanocapacitor surface and the neurons from *n* = 30 cross sections. (C) A FIB cross section of the marked region in (A) at the crossing of an Rx and Tx. The cell(s) have encompassed the electrode and small cavities have formed across the surface. (D) FIB-SEM cross sections of thin layer plasticized neurons (scale = 500 nm). i) This cross-section shows tight coupling between the cell(s) and the electrode as opposed to ii) where there is greater variability in the coupling.

**Figure 4:**
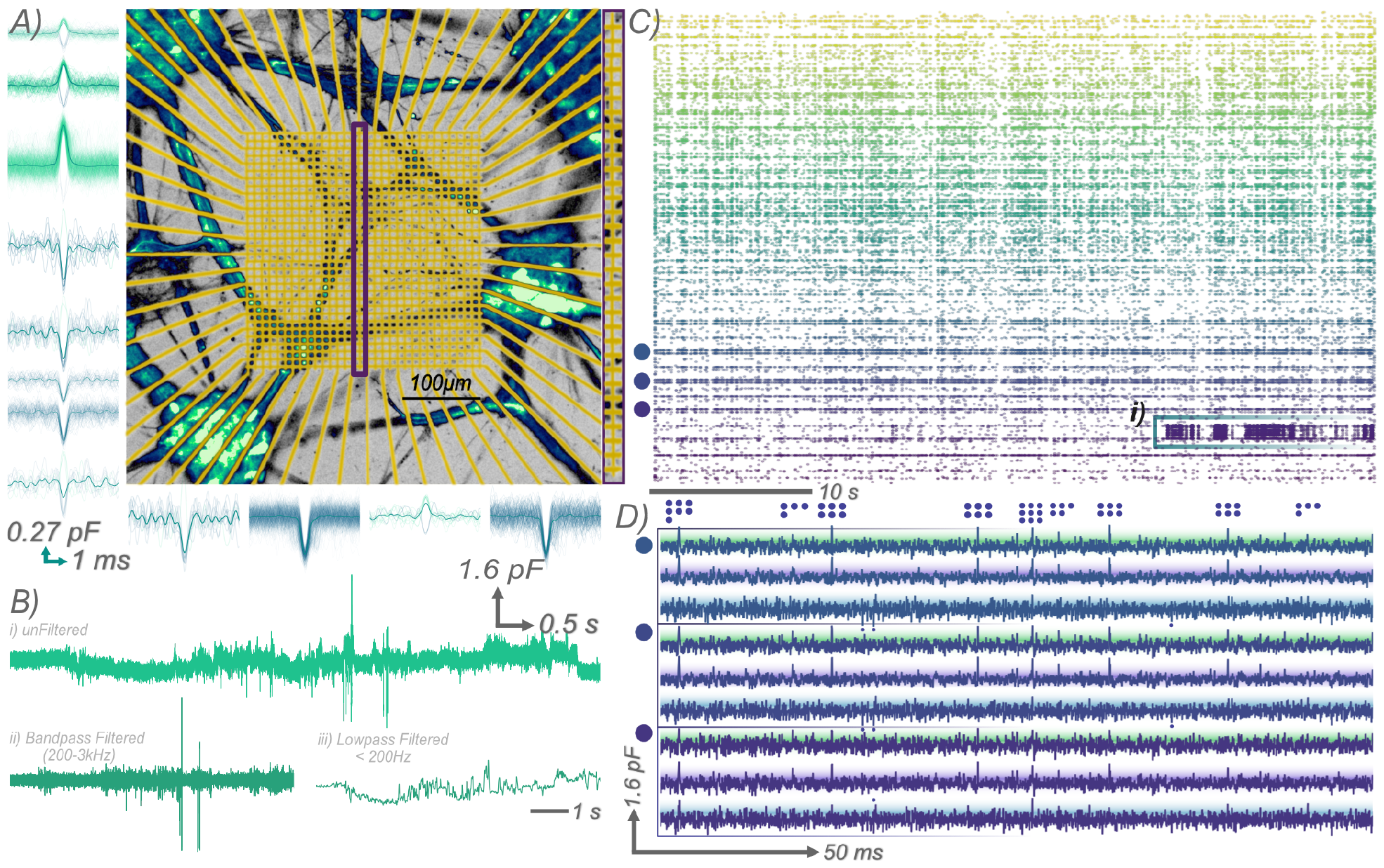
*In vitro* validation of 1024 parallel capacitive recordings of neural activity. (**A**) Primary cultures gCaMP8m stained at DIV14 on a 32×32 array of nanocapacitors. Electrodes and tracks are overlayed in yellow. Spike overlays of 12 electrodes from Rx17 during 40 s are the micrograph. (B) Un-, bandpass-, and lowpass-filtered traces. Low frequency changes are consistent with local field potentials, high frequency spikes with APs. (C) Raster plot of spikes detected on each electrode. Rx are sorted by color. i) Bursting activity is seen on one Rx. (D) Traces from 3 Rx (marked by dots) and 3 Tx (marked by color) showing common activity across electrodes. A selection of action potentials detected across electrodes is denoted above the topmost trace.

Neural recordings were then taken from 1,024 electrodes at 20 kHz with 9.7 μW/channel, comparable to systems of similar capabilities, but with a substantially smaller ASIC size of 5.26 mm^2^. Prior to electrophysiology, culture viability and activity were confirmed with calcium imaging (fig. S13). Positive, negative, and bi-phasic spikes were observed (fig. S14) consistent with conventional MEA recordings (*33*). Spikes were detected and overlayed for 12 of 32 electrodes associated with one Rx from a DIV14 culture consistent with microscopy images (Fig. 4A). Features of neural activity, such as low frequency changes resembling LFPs and single spike events, were captured (Fig. 4B). Figure 4C shows a raster plot of APs across all electrodes; bursting activity detected across a single Rx is highlighted. Correlated activity is expected for neighboring electrodes sharing a common input. We show activity for 3 Tx across 3 neighboring Rx (Fig. 4D). Activity is often detected across electrodes indicating one neuron coupling to multiple electrodes. Variation in electrodes detecting simultaneous activity indicates multiple neurons coupled to the sensing region.

## Conclusion

We demonstrated the first mutual capacitive sensors (both single and in 32×32 arrays) to monitor neural activity with comparable capabilities of conventional MEA systems (*34*), while enabling simultaneous sampling of 1,024 electrodes at 20kHz. Our approach harnesses code division multiplexing to reduce the electronics required to record from multiple electrodes in parallel. This reduction is proportional to the square root of the electrode number, as opposed to parity. However, integration of non-binary codes becomes imperative to expand the electrode count without increasing the carrier frequency or power consumption while retaining the current circuit implementation. Detailed investigation on electrode geometry, material, and isolation will be needed for specific applications to maximize signal-to-noise and minimize electrical crosstalk. Nevertheless, the adaptable fabrication process and ability to maintain robust signals even over extended trace lengths, renders nanocapacitor arrays highly amenable to stretchable/flexible probes (*35*–*37*) in contrast to conventional MEAs. Permissive fabrication, high channel count, and low power consumption overcome limitations in deploying HD-MEAs *in vivo;* therefore, capacitive MEAs have the potential to record brain wide single neuron activity and provide unprecedented insights into the functional origin of behavior.

## Acknowledgments

We would like to thank Dr. Anne Greet Bittermann of ScopeM for her scientific and technical assistance regarding the FIB-SEM. We would like to thank Gustavo Prack for his early contribution to the physical modeling. We would like to thank José Mateus for his input on early manuscript drafts.

## Author Contributions

Conceptualization: SW, TJ, AR

Investigation: SW, WC, YC, YV, KV, GA

Methodology: SW, AR, WC, YC, DK

Resources: AR, YV, TS

Visualization: SW, KV, YC

Funding Acquisition: JV, TJ

Supervision: JV, TJ, DM, NN, SW

Writing – original draft: SW, NN

Writing – review and editing: All

## Data and Material Availability

Data will be available from the ETH Research Collection. DOI: 10.3929/ethz-b-000660383

## Supplementary Materials

Materials and Methods

Fig. S1 – S16

References (38-42)

