## Supplemental Information for "Code Multiplexed Nanocapacitor Arrays for Scalable Neural Recordings"

### Main text Methods

*Biophysical neuron simulation:*

Finite element modelling of the soma of a neuron was done using COMSOL Multiphysics (Version 6.1.0.282) using the Electric Currents (EC) and Transport of Dilute Species (TDS) packages. The electric currents interface was used as the relaxation time of ions in solution is, for the distances involved, on a timescale similar to the generation of an action potential. The model constants were based on the electrodiffusive Pinsky-Rinzel model published by Sœtra et al (*1*). The only deviation from the published constants was that we did not weigh the diffusion coefficients in the extracellular vs. intracellular domains by the tortuosity. Infinite element domains were used to simulate boundary conditions at infinity to ensure local changes were captured. This specifically applied to ground and the concentration of each ionic species in contrast to compartmental models which often do not explicitly take distance into account. To ensure well established features in electrochemical systems could be captured, such as the formation of a double layer, boundary layers with a minimum thickness of 0.6 nm were used at all interfaces between domains. However, neither a quasistatic thermally induced potential (Boltzmann potential) nor potentials arising from specifically adsorbed molecules were included in the model. To improve solution stability, inconsistent stabilization of the dilute species physics was used with a coefficient $\delta_{id} = 0.05$ , which in essence adds a small perturbation to the diffusive flux preventing oscillations in convergence. Coupling of the potential solved by the EC to the TDS is natively supported by COMSOL, but coupling the charge density of the ions to the evolution of the electric field was done by adding the ionic charge density $\rho_{ion}$ to the total charge density:

$$\rho = \nabla \cdot D + \rho_{ion}$$

$$\rho_{ion} = \sum_{ion} z_{ion} c_{ion}$$

where $z_{ion}$ is the charge of an individual ion and $c_{ion}$ is the ionic concentration. Additionally, the standard derivation of the membrane potential in HH neuron models relies on the Nernst equation which is strictly valid only under equilibrium conditions. As such, it was necessary to derive the membrane potential from the ionic flux and reported values for what is commonly called membrane capacitance $C_m$:

$$\frac{C_m}{F}\frac{\partial V_m}{\partial t} = J_{Na} - J_K + J_{Cl} - 2J_{Ca} + K_{stim}$$

Where $J_x$ is the total ionic flux for a given species, $K_{stim}$ is an additional potassium flux used to induce action potentials, and $F$ is the faraday constant. Ionic reversal potentials, HH formalism, and passive and active membrane conductances remain the same as those in the Pinsky-Rinzel model. A time dependent study for 5 induced APs across 15ms with reported values every 50μs was run on an ARM based desktop computer with 10 CPU cores with the average simulation taking 12hrs to complete. Relevant data from each simulation was then exported and evaluated in Python.

*ASIC and PCB design:*

The theoretical basis for circuit function was validated mathematically using MATLAB and Cadence Virtuoso with optimization for low power consumption and noise. The circuit was designed in Cadence Virtuoso and manufactured in 180 nm CMOS technology (TSMC). After manufacture, electrical testing of chip performance was compared with the design specifications. The motherboard and daughter card for the ASIC were designed in Altium.

*MEA fabrication:*

A borosilicate wafer (MicroChemicals) was Oxygen plasma cleaned for 2 minutes prior to vapor deposition of a 5nm Ti adhesion layer followed by 150 nm of Au and a final layer 5nm of Ti as an adhesion layer for subsequent steps (Plassys). After a 5-minute bake at 200˚C, the wafer was placed in hexamethyldisilazane (HDMS) atmosphere to improve subsequent photoresist adhesion. Next, a photoresist layer (AZ1518, MicroChemicals) was spin coated (40 s at 4 krpm), soft baked (90 s at 110°C), exposed (EVG 620, 80 mJ), developed (40 s in 4:1 AZ 400), and hard baked (60 s at 115°C) to act as an etch mask. The top Ti adhesion layer was removed using reactive ion etching (RIE, Oxford Instruments, NGP80), Au was then chemically etched (potassium iodine (KI)), and the photoresist mask layer stripped. After ensuring no obvious photoresist residues remained, the wafer was oxygen plasma cleaned for 2 minutes and 200 nm of SiO2 was vapor deposited on the surface. This was followed by deposition of another 5 nm Ti adhesion layer followed by 170 nm Au. Photoresist patterning and subsequent etching proceed the same as the first layer of Au structures. Next, the SiO2 layer was RIE etched with the top Au structures acting as an etch mask to preserve the insulation between the overlapping areas of the top and bottom layers.

*MEA packaging:*
A PCB was designed using DipTrace to act as an interface between the MEA and the daughtercard. Wafers containing the MEAs were diced and each sample inspected for defects. Conductive bonding between the MEA and the PCB was done by manually placing a drop of silver epoxy (MG Chemicals, 8331) on the pads of both the PCB and MEA, aligning the two, pressing them together and baking overnight at 80˚C. A polydimethylsiloxane (PDMS 184, Dow Chemicals) or epoxy (EPO-TEK 353ND-T) ring and coating were placed around the bonding region to prevent leakage of neurotoxic compounds from the PCB to the culture medium.

*Electrode response to concentration changes:*
The response of the MEA to changes in concentration was evaluated by using the ASIC to compare the output of a capacitive pair to a solid-state capacitor. The relationship of the capacitance to the osmolarity was found by starting with diluted (0.9x) neurobasal culture medium doing serial additions of 1.5x PBS. The transient response of the chip to sudden changes in concentration was performed by pipetting 1.5x PBS in decreasing volumes of MilliQ water.

*Cell culture:*
Animal experiments were approved by the Cantonal Veterinary Office Zurich. Primary neurons were taken from E18 Sprague-Dawley rats and dissociated according to established protocols (*2*). Briefly, dissected tissue was digested in papain for 15 minutes followed by subsequent washes with neurobasal medium, trituration, and cell counting (Countess 1, ThermoFisher). Substrates were prepared for cell culture by removing debris and contaminants through ultrasonication in 2% SDS followed by two minutes of oxygen plasma cleaning. Cultures were performed without additional surface coating. Hippocampal cells were seeded at a density of approximately 100k cells/$cm^2$. Culture medium was changed the day post seeding and twice weekly afterwards.

*FIB-SEM:*
Samples were fixed in EM-grade glutaraldehyde in sodium cacodylate buffer then washed 3 times with buffer followed by post fixation in 1% $OsO_4$. After washing with water, samples were stained with 1% uranyl acetate, washed, and serially dehydrated in ethanol. Serial embedding with Epon resin was done followed by either thin layer plastification (TLP) (*3*) or ultra-thin resin embedding (UTP) (*4*). For TLP the embedded samples were either centrifuged in an Eppendorf or blotted with filter paper. For UTP, the samples were rinsed with 3 mL of water free ethanol 30 times. Samples were then placed upright in an oven at 60˚C for 2 days to allow for polymerization. Samples were then mounted on SEM stubs and sputtercoated with 4 nm Pt/Pd. Imaging and milling were done using a Helios Nanolab 600i (FEI). Overview images were taken at 30kV and 0.17nA with backscatter electron detection to identify regions of interest. The sample

was tiled 52˚ for perpendicular ion milling and a front trench was cut with Ga ions at 30kV and 9nA. Subsequent cross sections were cut with Ga ions at 30kV and 2.3nA and imaged by SEM with an in-lens scattered electron detector in back scatter mode at 2kV and 1.4 nA. Tilt correction for perspective distortion of the closeups of the cross sections was performed.

*Analysis of Neuron/Electrode Interface:*
Images were manually preprocessed to isolate both the regions of interest and label the substrate (*i.e.*, the electrode and isolation material). Edge detection based on the first and second derivative of the pixel intensity was done with manual thresholds used for each image. A simple heuristic was used to identify the first edge between cell material and the extracellular space. Edge detection scripts were written in python.

*Neuron Recordings:*
Recordings were taken between DIV9 and DIV14 at 15-minute intervals at ambient conditions in an electrically shielded cage. Three 6-layer PCBs were used: a breakout board for interfacing with the ASIC with the peripheral circuitry to allow the IC to function, a daughter board with the ASIC, and the MEA PCB for interfacing between the breakout board and the electrodes (fig. S10). Orthogonal codes were generated off chip by a controller and passed to the ASIC drivers. Four 8 channel logic analyzers (Selee) collect the output of the chip and pass the data to computers for subsequent analysis. Due to memory constraints individual recordings were limited to between 30 and 40 s. Binary data from the logic analyzers is passed to custom MATLAB scripts which convert the raw data to the relative capacitance changes for each electrode.

*Analysis of Neural Recordings:*
Custom Python 3.6 scripts were written to analyze the electrophysiology data. For analysis involving action potentials, a second order bidirectional bandpass filter using second order sections with a passband of 100 - 5kHz was implemented. Potential spikes were found by identifying all datapoints with a magnitude greater than 4.5$\sigma$. If multiple datapoints within a 2ms fell above this threshold only one point was retained. For spike overlays from each electrode, a 4ms window around the spike time, the maximum magnitude was found and the time shifted so this value was considered 0.

**Methods for Supplementary Figures:**

*Modeling Applied Potentials in Solution:*
Finite element modelling of the nanocapacitors and the effect on the solution of an applied potential were done using COMSOL Multiphysics (Version 6.1.0.282) using the EC and TDS packages. Diffusion coefficients and model parameters were the same as those used in the biophysical model of a neuron. Three capacitors sharing a common condenser plate (3 Tx, 1 Rx) were included in the model. A factorial study of the effect of potential and frequency on the solution was performed. The potentials used were 15 µV to 12mV, 24 mV, 36 mV, 48 mV, 60 mV. The frequencies used were 1 kHz, 10 kHz, 50 kHz, 100 kHz, 500 kHz, 640 kHz, and 1 MHz. Geometry of the capacitors modeled follows the nominal values for the capacitor design (Fig. 2B).

*Calcium Imaging:*
Neuronal cultures were transduced at DIV5 with an AAV produced by the UZH Viral Vector Facility to express GCaMP8m under the hsyn1 promoter. Titer volume was adjusted to have approximately 10k viral particles per cell. Imaging was done in incubator conditions (5% $CO_2$, 37˚C) using a confocal laser scanning microscope (Olympus FluoView3000) using a resonant scanner at a framerate of 30 fps. Activity was monitored across 10 minutes, though recordings of ~1 minute were taken. Imaging was performed directly prior to electrophysiology; between DIV9 and DIV14.

*Electrochemical Impedance Spectroscopy:*

Electrode impedance measurements were taken in 1x PBS from frequencies between 1kHz and 800kHz with a probe potential of 10mV using a semiconductor device analyzer (Agilent B1500A). Measurements were taken between an Rx and a Tx with all other traces left floating.

*Preliminary Electrophysiology:*
Experiments validating capacitive recordings of neural activity using CDM were performed using a PCB level implementation of the circuit with planar electrodes fabricated using standard liftoff techniques. Experiments were performed at 37˚C using $CO_2$ independent medium (Hibernate E) on DIV24. The capacitance of the MEA was compared to a solid-state capacitor rather than the total capacitance of the entire Rx line. Calcium imaging was performed prior to recordings to ensure cultures were active.

**Methods References**

(1) Id, M. J. S.; Id, G. T. E.; Id, G. H. *An Electrodiffusive , Ion Conserving Pinsky- Rinzel Model with Homeostatic Mechanisms*; 2020.
(2) Mateus, J. C.; Weaver, S.; Van Swaay, D.; Renz, A. F.; Hengsteler, J.; Aguiar, P.; Vörös, J. Nanoscale Patterning of in Vitro Neuronal Circuits. *ACS Nano* **2022**, *16* (4), 5731–5742.
(3) Kizilyaprak, C.; Bittermann, A. G.; Daraspe, J.; Humbel, B. *Electron Microscopy*; Kuo, J., Ed.; Methods in Molecular Biology; Humana Press: Totowa, NJ, 2014; Vol. 1117.
(4) Belu, A.; Schnitker, J.; Bertazzo, S.; Neumann, E.; Mayer, D.; Offenhäusser, A.; Santoro, F. Ultra-Thin Resin Embedding Method for Scanning Electron Microscopy of Individual Cells on High and Low Aspect Ratio 3D Nanostructures. *J. Microsc.* **2016**, *263* (1), 78–86.


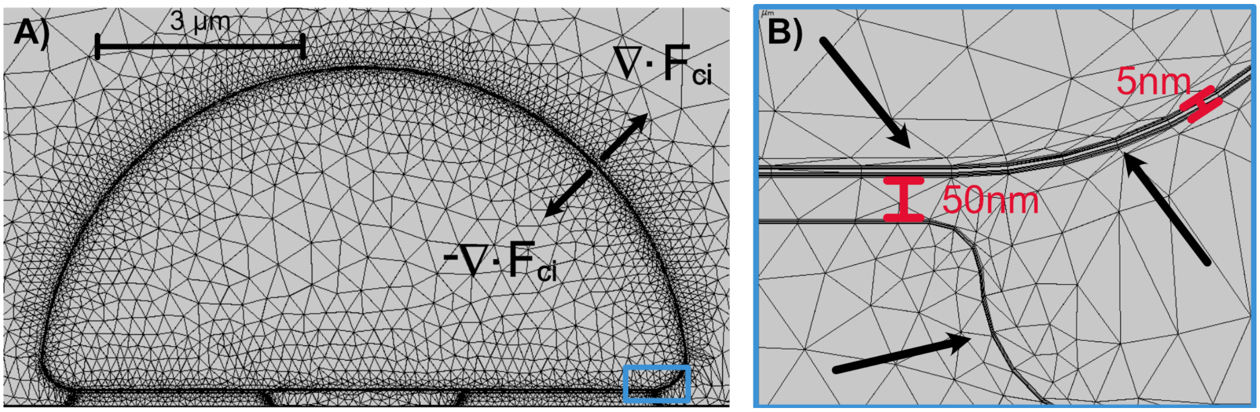


**Supplementary Figure 1: Finite element model of neural activity on two electrodes**. **(A)** The neuron is modeled as a hemisphere with a diameter of 10 µm sitting above two 200 nm thick gold electrodes. The flux across the membrane, $F_{ci}$, uses the Hodgkin-Huxley equations to govern the permeability across the membrane. (B) The thickness of the cell wall was set to 5 nm. The cleft height in this image was 50 nm. Six boundary layers (denoted by arrows) with a thickness ~1 nm were included in the mesh for each boundary contacting the ionic solution to allow for double layer formation. All edges were rounded to prevent singularities in the solution of the electric field.

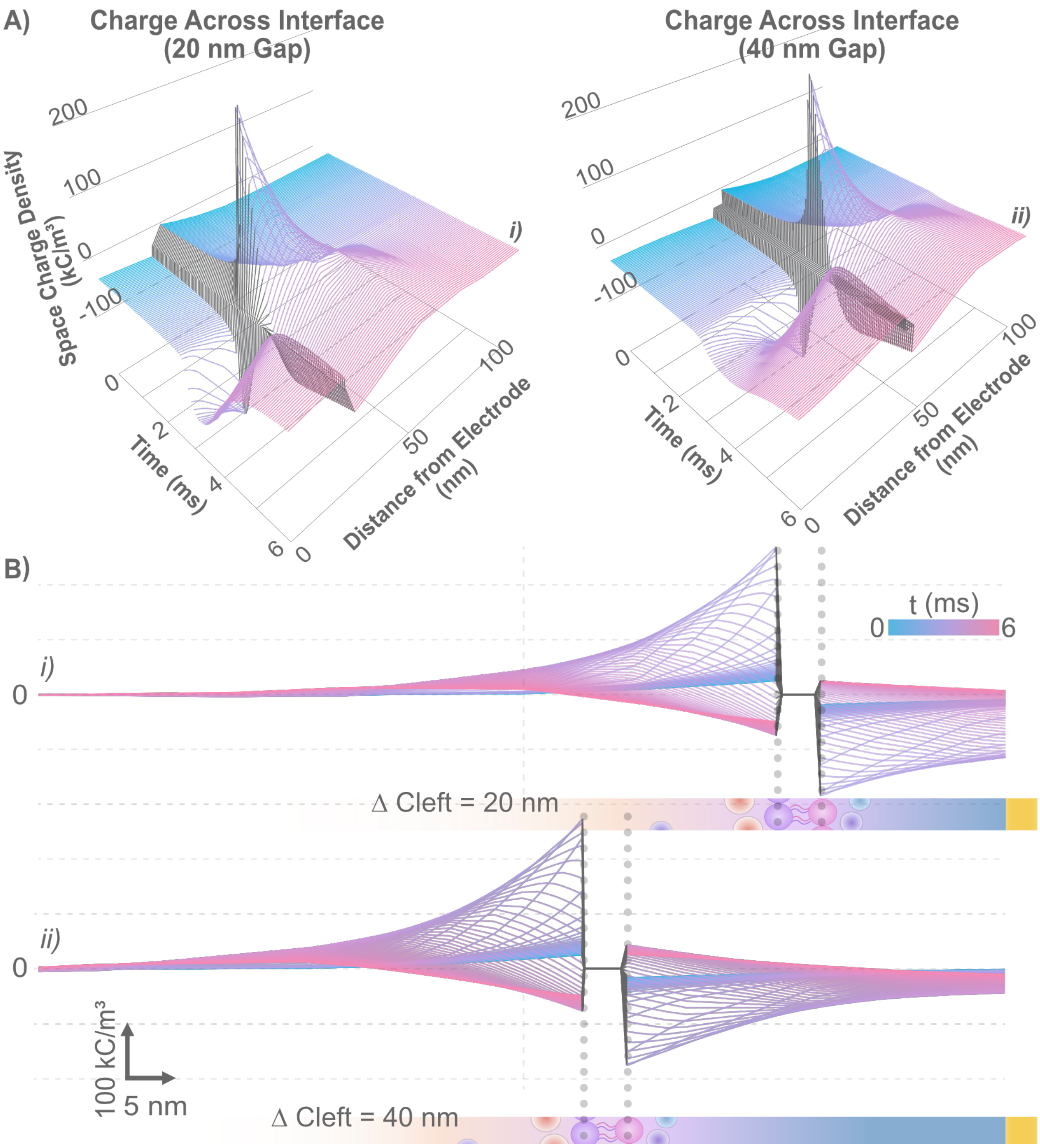


**Supplementary Figure 2: (A)** Charge density across the gap between the electrode and cell membrane (cleft) for i) a 20 nm cleft height and ii) 40 nm cleft height for a single AP. (B) 2D projection of the same data for i) 20 nm cleft and ii) 40 nm cleft. The charge density at the inner and outer surface of the cell membrane changes minimally with changes in cleft height; however, the charge density at the electrode surface differs by ~3x (-40kC/$m^3$ vs -115kC/$m^3$). Additionally, for a 20 nm separation the charge across the cleft does not change substantially at the peak of the AP whereas there is a noticeable decrease for a 40 nm gap. 100kC/$m^3$ is equivalent to unbalanced charge of 1mM for a monovalent ion.

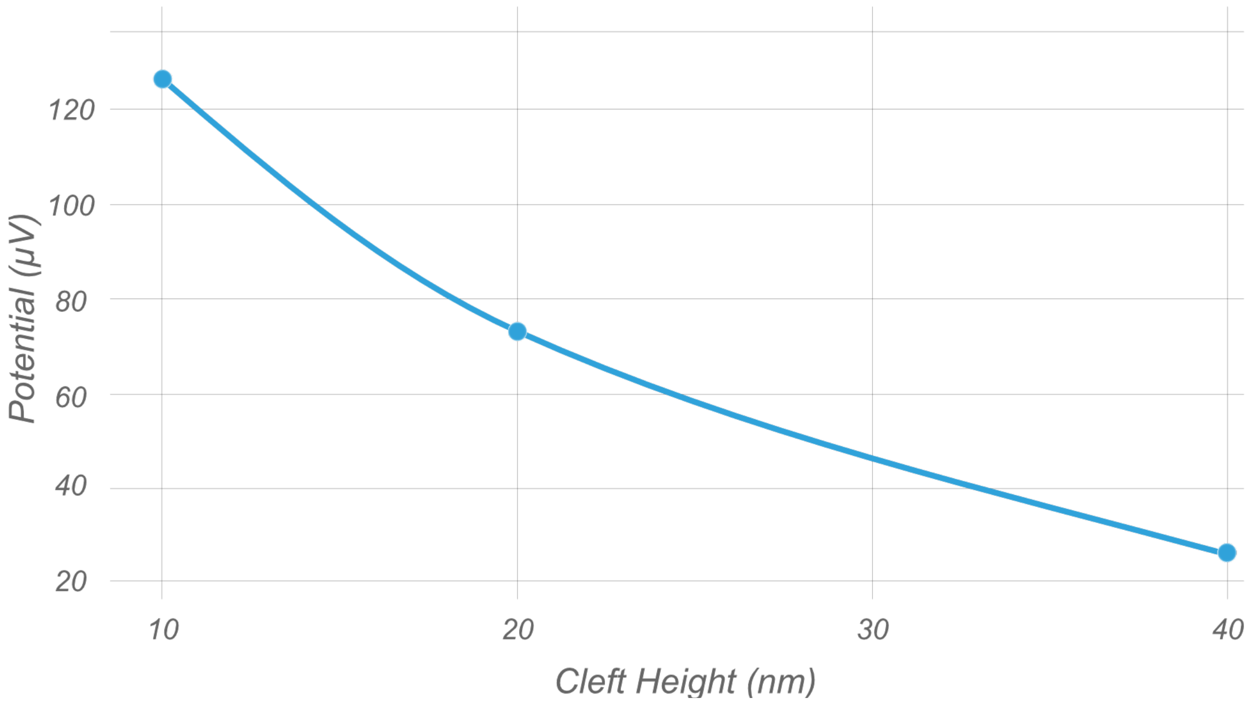


**Supplementary Figure 3: Electrode potential variation with cleft height.** Maximum potential at the electrode during a series of 5 action potentials versus the distance of the cell membrane to the electrode surface. In our 2D simulation, which is symmetric, considering cleft height is appropriate; however, in physical systems this is more appropriately the cleft volume.

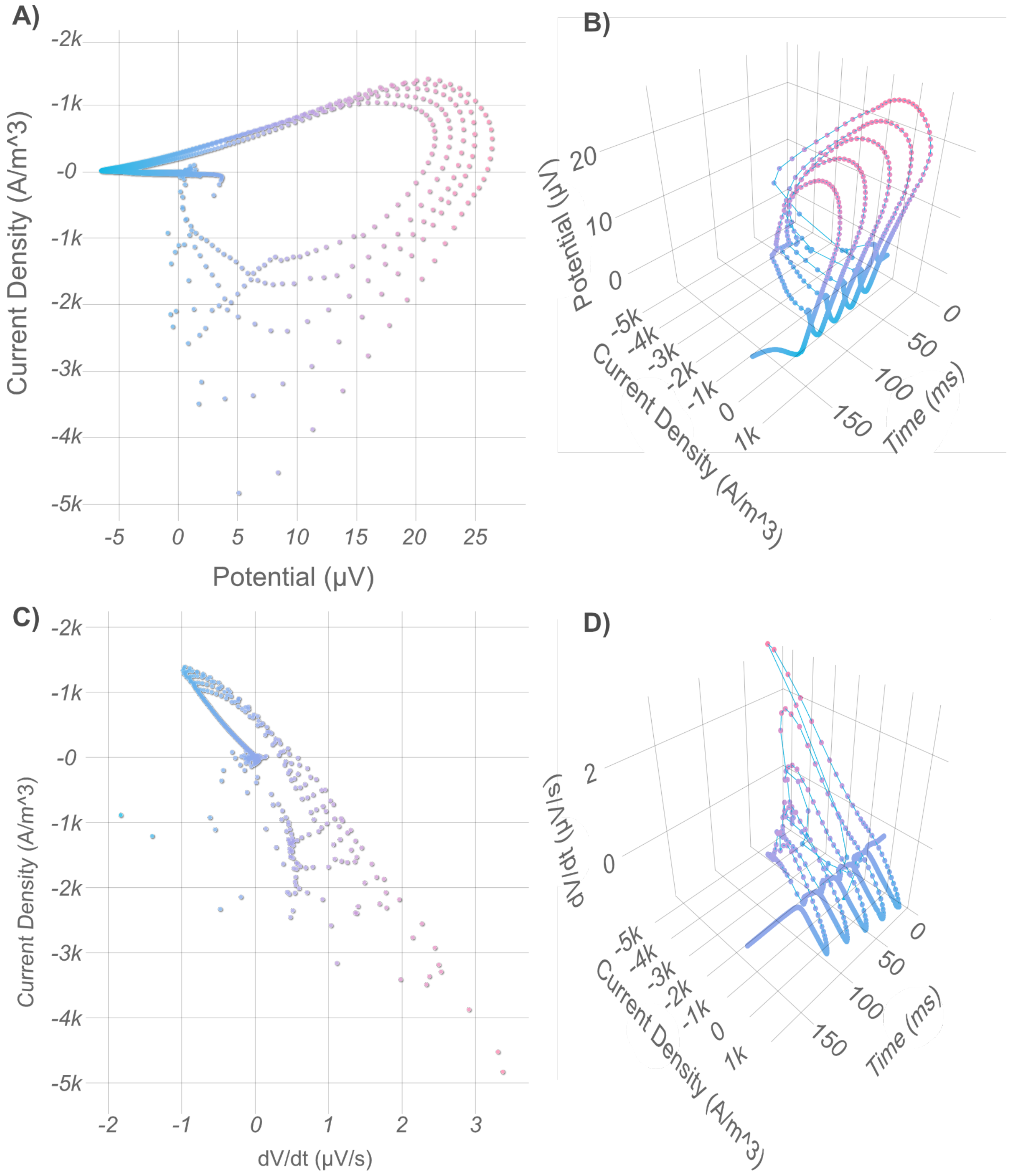


**Supplementary Figure 4**: **Simulation results for differential capacitance of the electrode for 5 action potentials.** (**A**) Current density vs. potential. (B) The same data as A, but including plotting versus time. (C) Current density vs. dV/dt. The slope at each point corresponds to the differential capacitance. (D) The same data as in C, but plotted versus time.

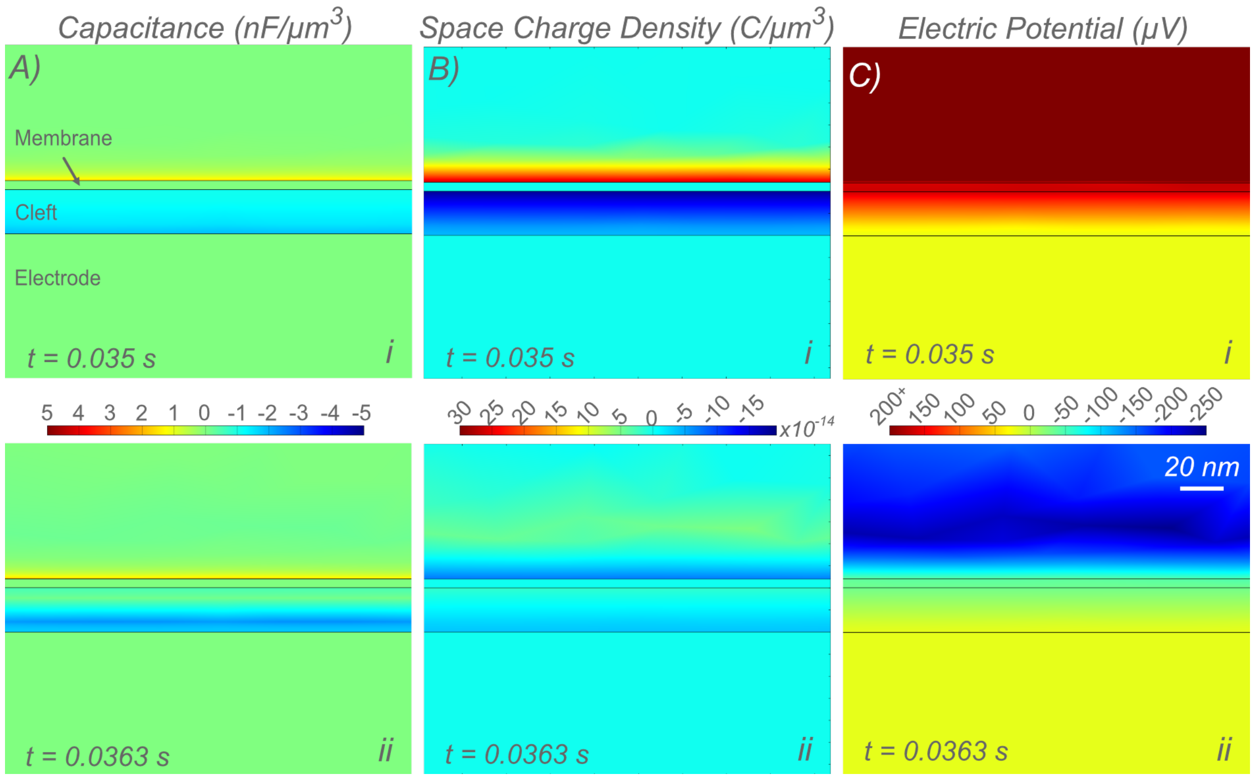


**Supplementary Figure 5: Variation in the capacitance, charge, and potential across the cleft during an action potential. (A)** The capacitance ($\rho/\phi$) across the cleft (20 nm) at i) the rising and ii) the falling edge of an AP at equipotential points at the electrode (20mV). In addition to differences at the electrode surface as shown in Fig. 1E (for 40 nm cleft), we see changes in capacitance across the cleft. The capacitive contribution to the detected signal of each volume element will depend on the cell coupling, geometry, frequency, and recording technique (*e.g.* transimpedance *vs.* potential coupling); however, the largest contribution to the detected signal should come from the volume elements closer to the electrode surface. Conduction of ions in solution adds to dielectric loss, which is applicable in time dependent systems. Loss exceeding the real component of the permittivity would result in negative capacitance. This arises in the instantaneous calculation of capacitance for elements within the cleft due to the current contribution to the electric field (*ergo* potential). (B) Shows the corresponding charge density and (C) the potential.

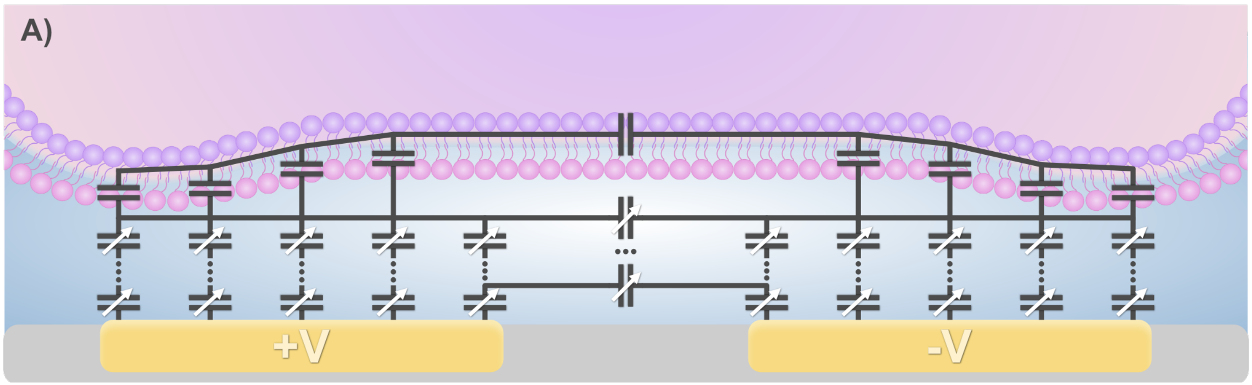


**Supplementary Figure 6: Electrical circuit approximation of capacitance changes in the cleft at high frequencies. (A)** Unlike conventional circuit models of the neuron/electrode interface which aim to approximate ionic flux governed by Nernst-Plank-Poisson equations and the subsequent ionic coupling to the electrode, the distributed capacitor network depicted describes the coupling between the condenser plates. The variable capacitance of each element within the cleft depends both on neural activity and relative location to the cell and each condenser plate. While not strictly accurate due to charge transfer through ion channels, the cell membrane is assumed to have constant capacitance as the volume occupied by bound lipids with low polarizability is in excess of the locations with conduction. At low frequencies, the contribution of the applied potential to the charge distribution within the cleft cannot be neglected.

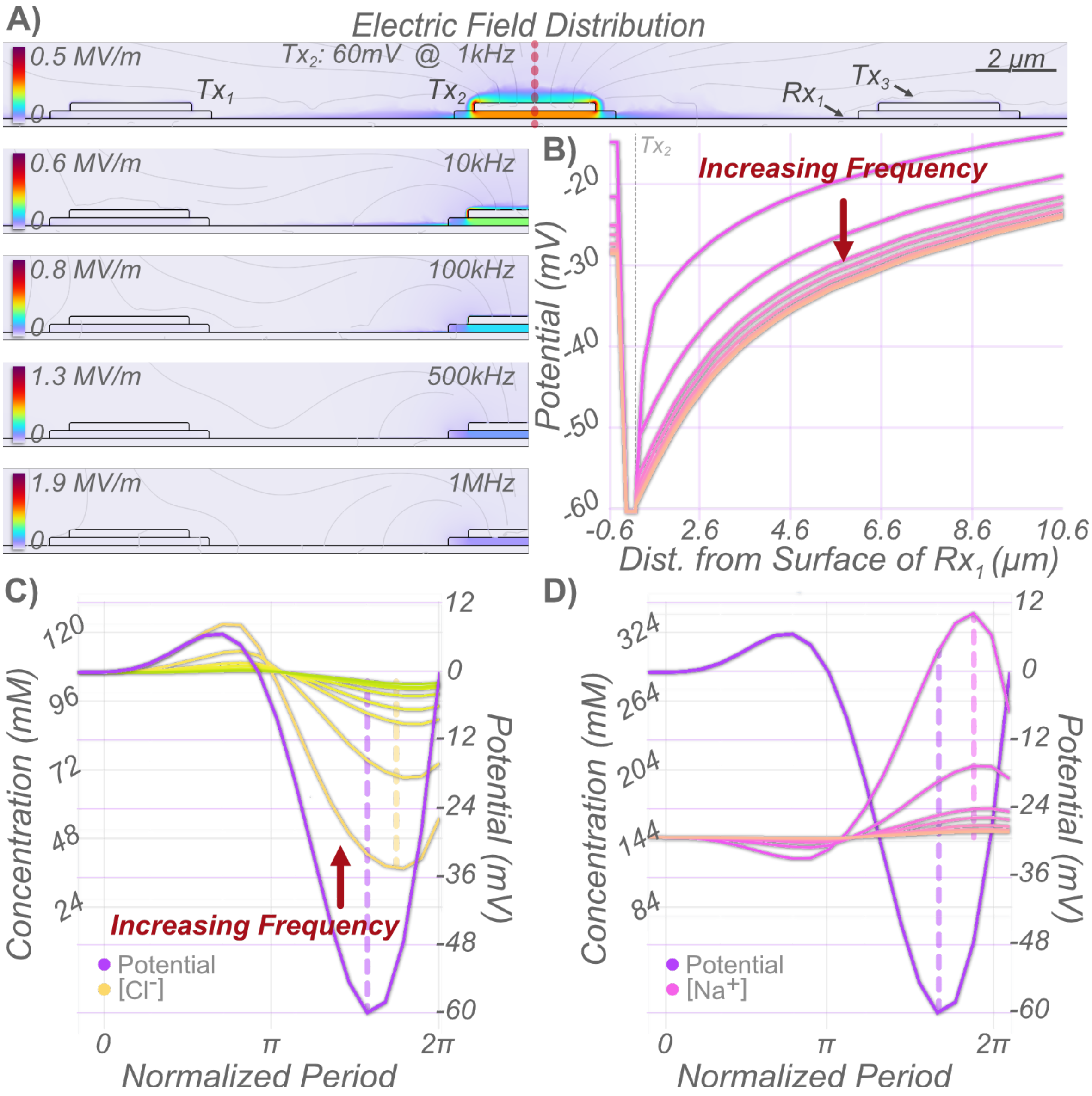


**Supplementary Figure 7**: **Frequency response to applied potential of nanocapacitors in simulation.** (**A**) Three capacitors ($Tx_{1\text{-}3}$) with a common condensor plate ($Rx_1$) with the dimensions outlined in Fig. 2B were simulated and the penetration of the electric field into the solution when a 60mV potential was applied to $Tx_2$at 1kHz, 10kHz, 100kHz, 500kHz, and 1MHz to investigate the impact of an applied potential on the ionic solution. The potential of the other electrodes ($Tx_1$, $Tx_3$, $Rx_1$) was left floating. As the frequency increases the relative intensity of the electric field becomes more restricted to the surface of $Tx_2$ and in the dielectric due to decreased ion conductance in solution. The dotted red line denotes where data for subsequent graphs was taken. (B) The maximum potential from $Rx_1$, through the dielectric, to 10.6µm into the solution for each frequency represented in A. As the frequency increases, the potential extends deeper into the solution. (C) Variation of the $Cl^-$ concentration and potential at the electrode surface at different frequencies. Time was normalized to the period of the applied potential. As the frequency increased, the change in concentration at the electrode surface was reduced. At 1Mhz the magnitude of the maximal change in concentration is 4mM. (D) The same as C, but for the concentration of $Na^+$. At 1Mhz the magnitude of the maximal change in concentration is 5mM. The maximum magnitude of the potential and of the concentration are offset by ~12% of the period for 1kHz. This offset increases as the frequency increases.

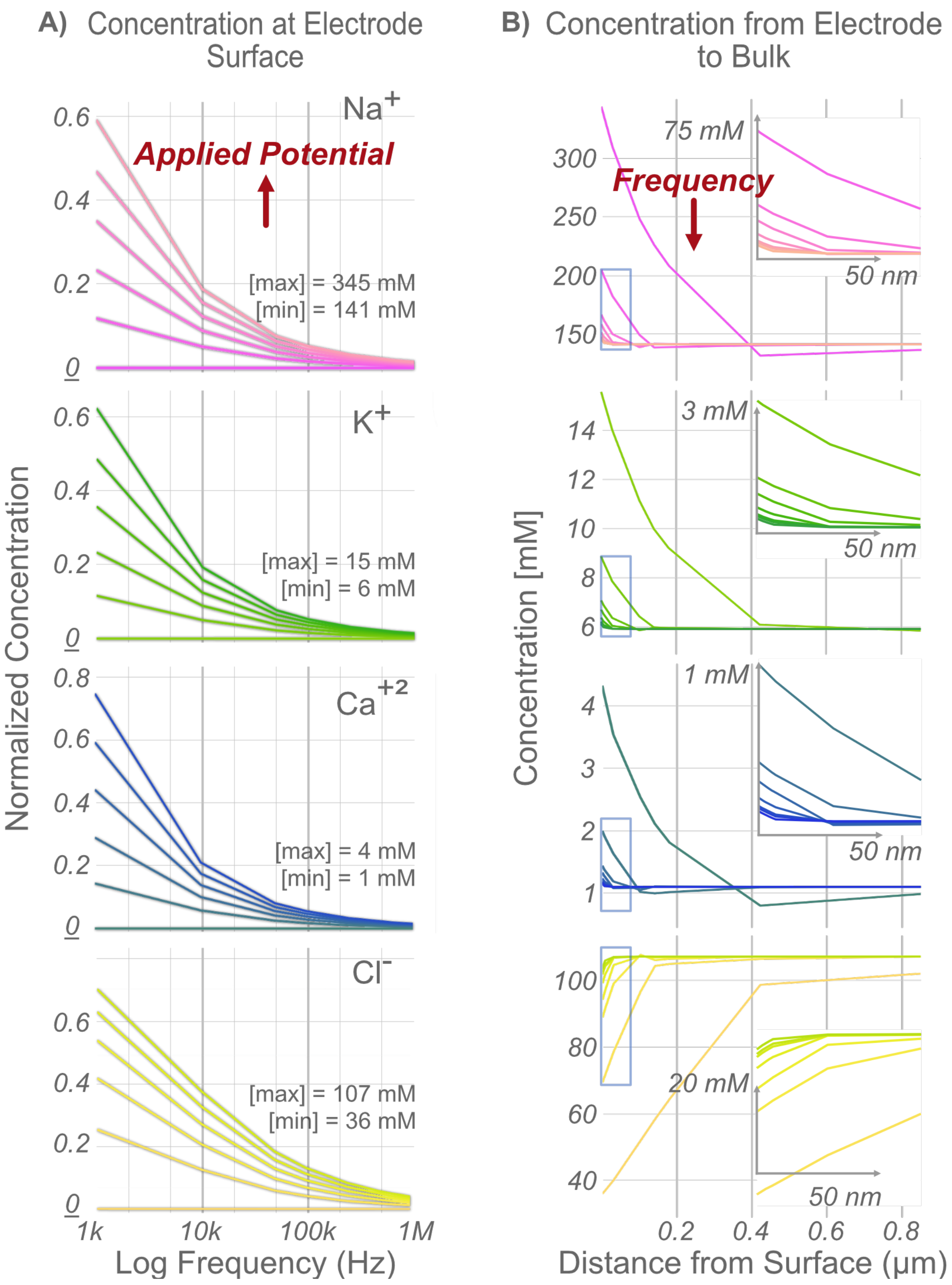

**Supplementary Figure 8**: **Concentration of each species across potential and frequency in nanocapacitor simulations.** Data for this figure comes from the same simulation as fig. S6 **(A)** The maximum concentration at the electrode surface for each applied potential vs. frequency. Normalization is division by the maximum concentration then subtracting the minimum/maximum. As the applied potential increases from 15 µV to 12mV, 24 mV, 36 mV, 48 mV, 60 mV the concentration at the surface increases. While the trend is maintained, increase in frequency reduces impact of the potential. (B) Variation in concentration of each species from the electrode surface for an applied potential of 60 mV for 1 kHz, 10 kHz, 50 kHz, 100 kHz, 500 kHz, 640 kHz, and 1 MHz. As the frequency increases, the difference in concentration between the electrode surface from the bulk decrease in magnitude and return to the bulk concentration across a shorter distance. Insets show the area highlighted by the blue box.

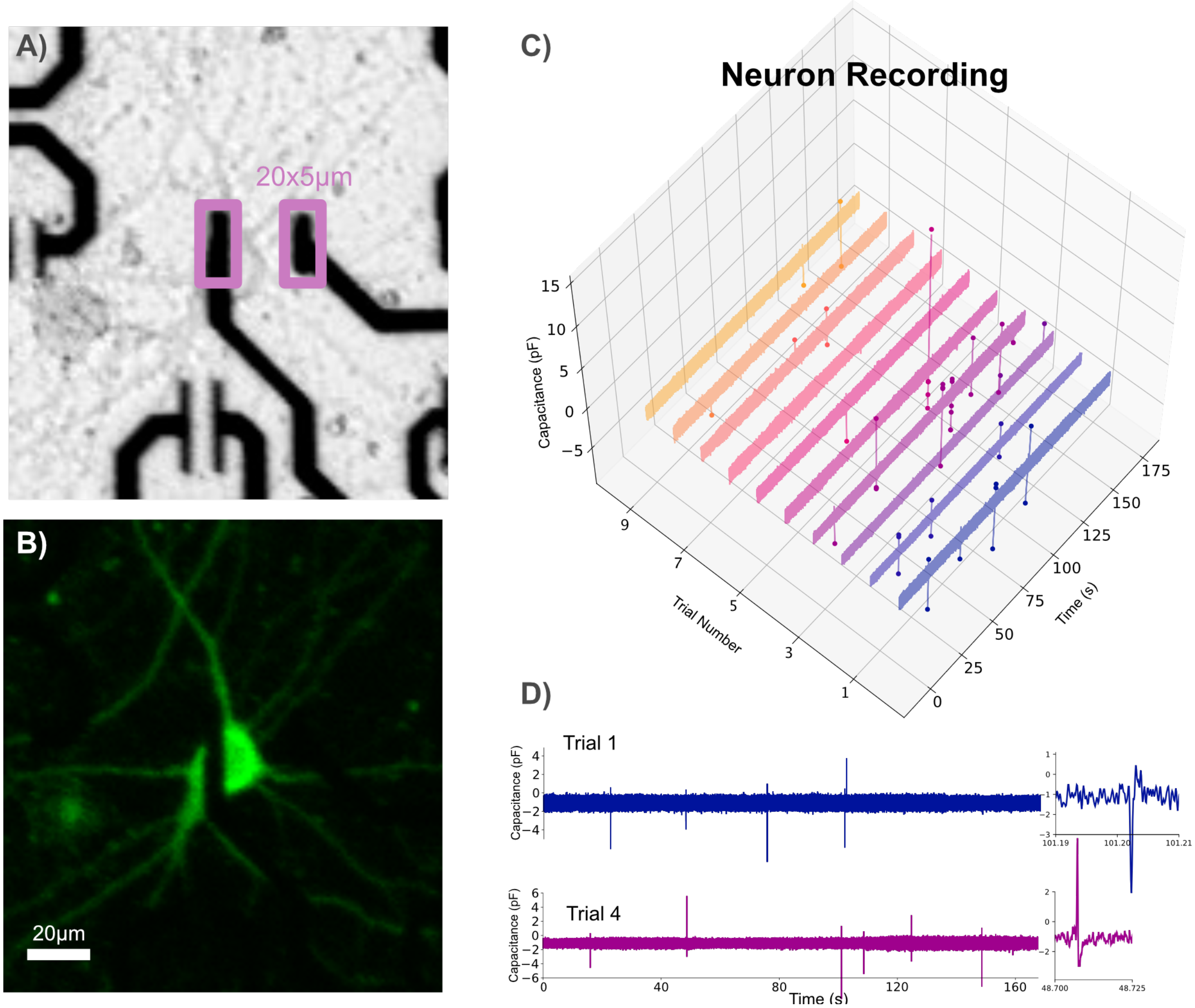


**Supplementary Figure 9: PCB level tests of code division multiplexed capacitive recordings. (A)** Brightfield image of a neuron at DIV24 on top of a coplanar capacitor. B) the same as in A, but showing GCaMP6m expression. C) Electrical recording of the neuron pictured in A across ten 3 minute recordings. D) Traces from trials 1 and 4 showing the detection of action potentials. Insets show closeup of action potentials.

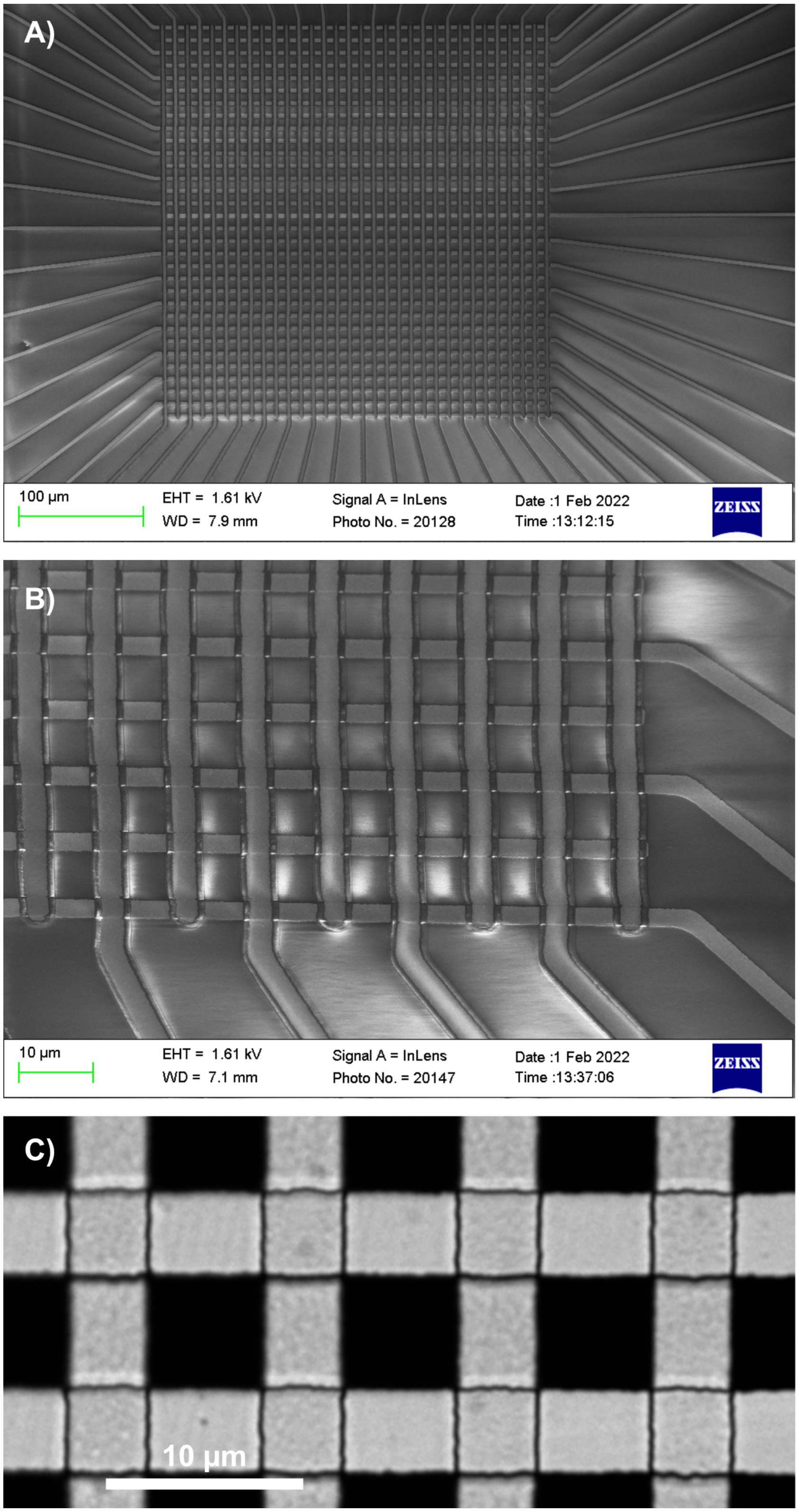


**Supplementary Figure 10: Example images of the nanocapacitor array. (A)** SEM overview of the array directly post fabrication. (B) Zoomed in image of the bottom right corner from the array shown in A. (C) 20x brightfield image of a different nanocapacitor array.

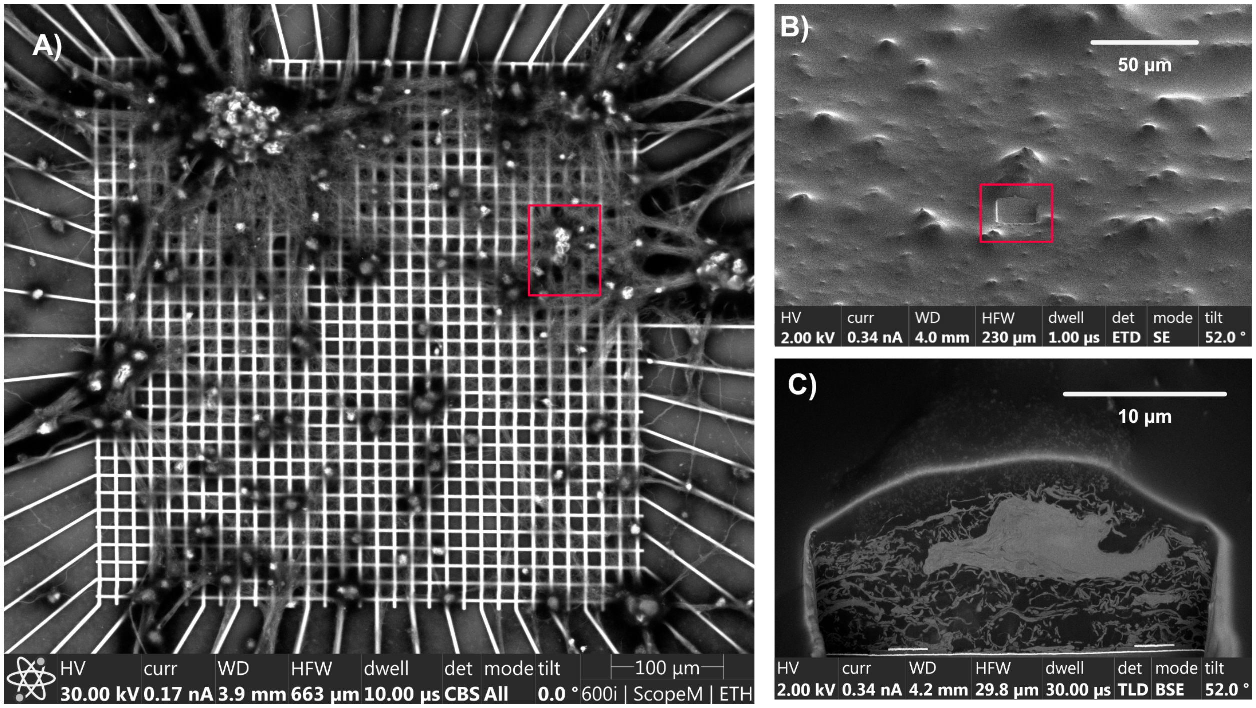


**Supplementary Figure 11: FIB-SEM of TLP fixed 32x32 electrode array. (A)** Overview image of neurons growing on a 32x23 capacitor array. (B) Initial FIB cut of the region marked in red in A. (C) Cross section of neurons and neurites growing on the array. Segments of the interface between the electrode and the neuron were used for analysis of the cell/electrode interface.

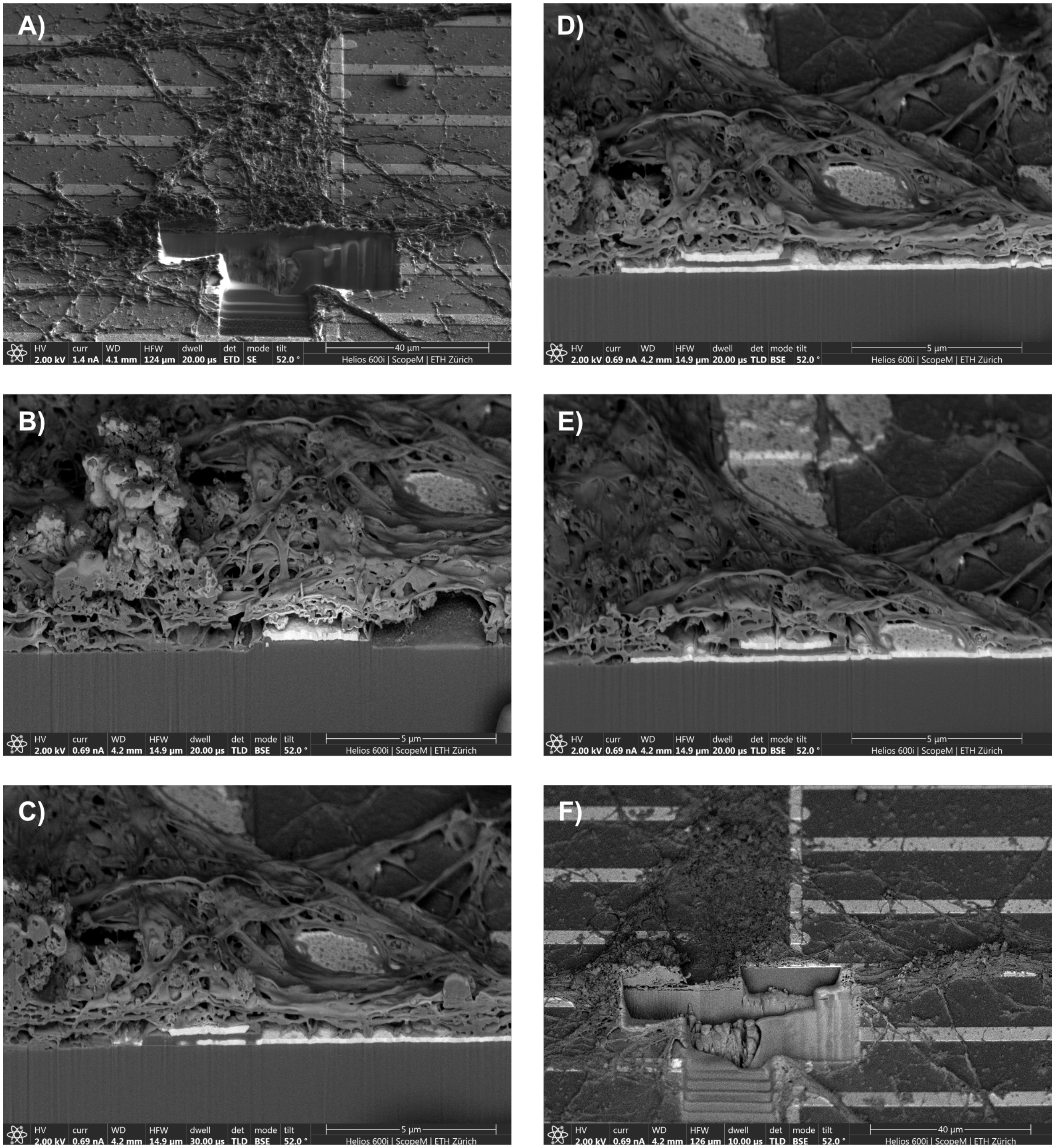


**Supplementary Figure 12: Sequential images of FIB cuts from a UTP fixed culture on a 32x1 electrode array. (A)** Overview image of the region of interest after initial FIB cuts have been made. (B-E) Sequential images of the FIB cuts. (F) Overview of the region of interest after all FIB cuts have been made.

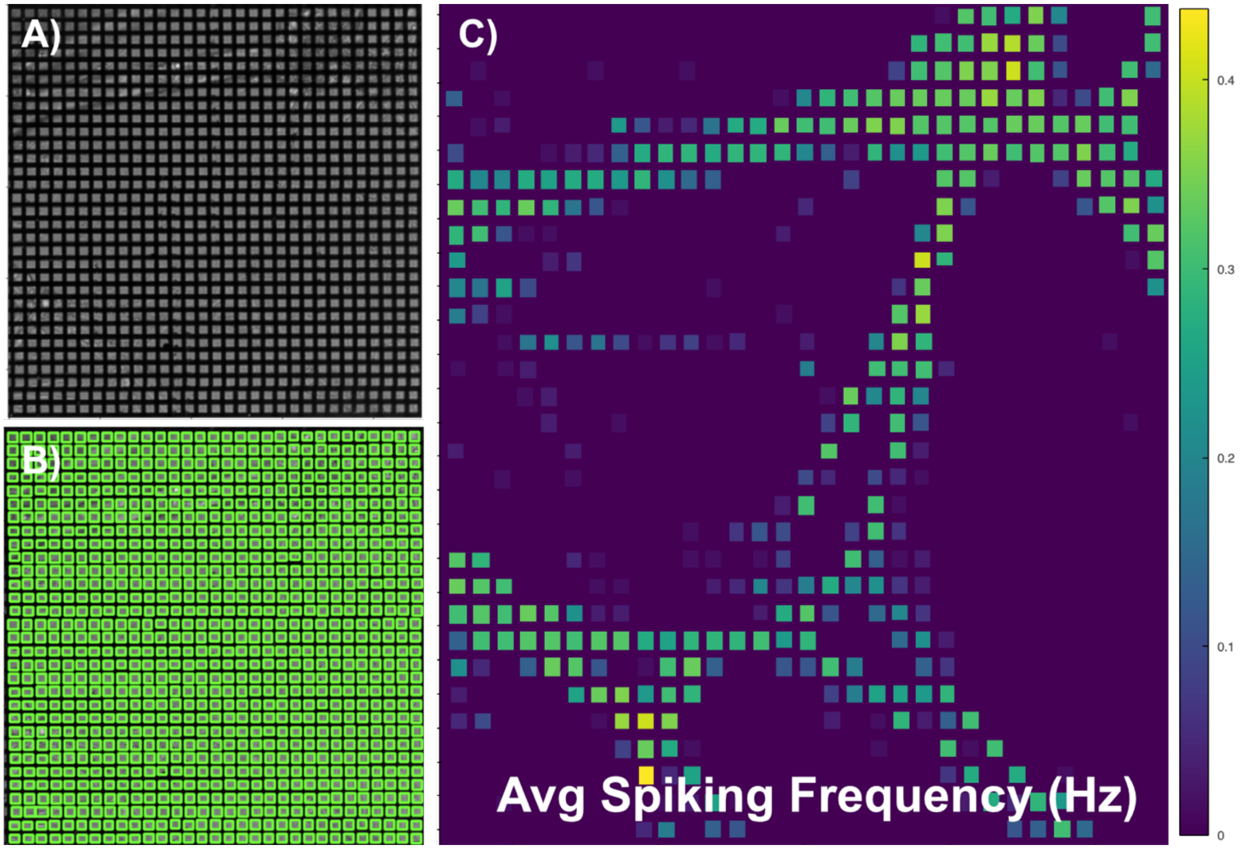

**Supplementary Figure 13: Verification of culture activity using calcium imaging. (A**) Brightfield image of the MEA. B) Segmentation of the calcium imaging into ROI between the Rx and Tx. C) Average spiking frequency of each ROI

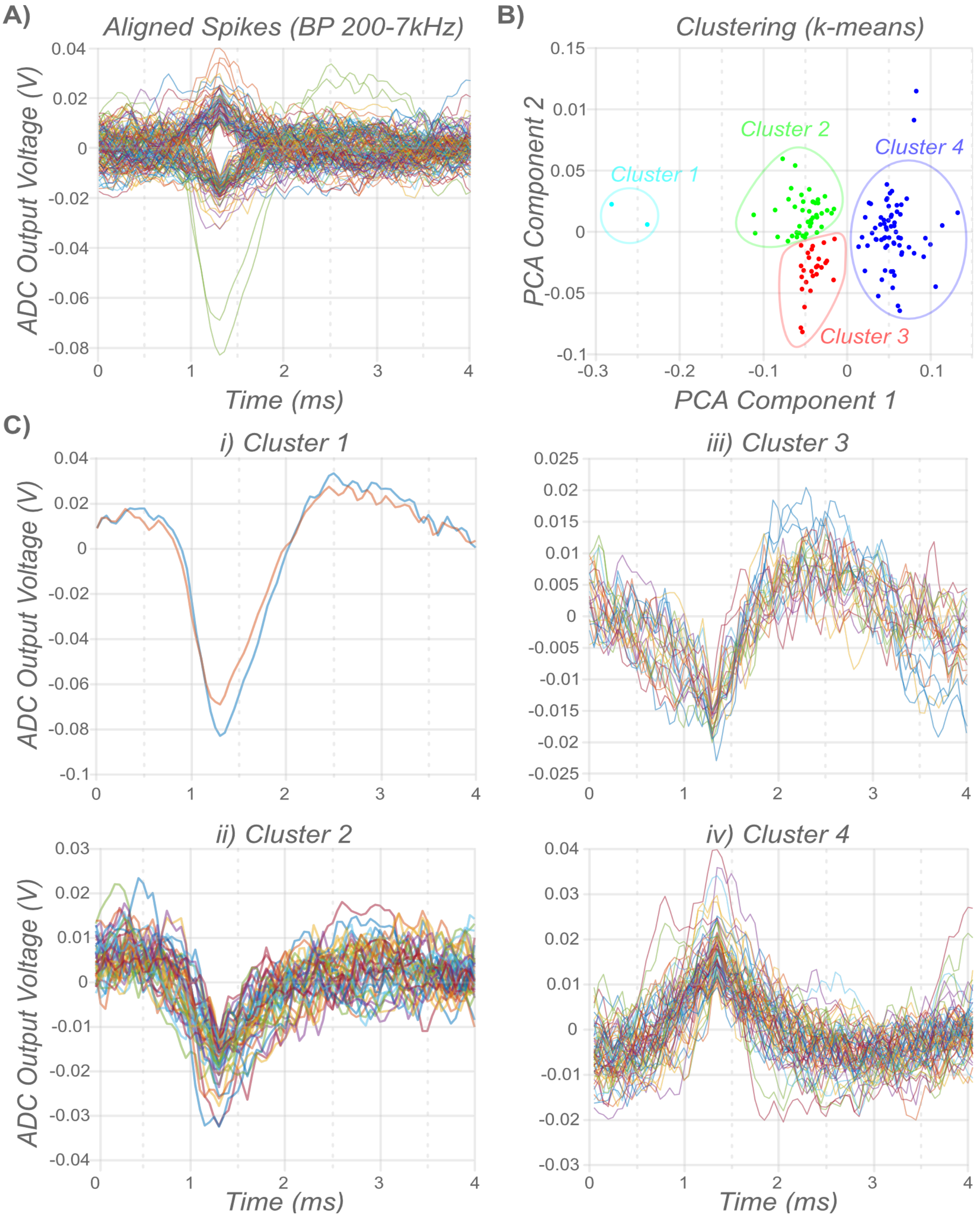


**Supplementary Figure 14: Variation of spike shapes from a single RX. (A)** Overlay of all spikes detected on a single RX aligned to the maximum magnitude of the spike. (B) Results of clustering using the K-means algorithm. There are 4 clusters found, though clusters 2 and 3 may not be separate. (C) Overlay of the spikes from each of the clusters.

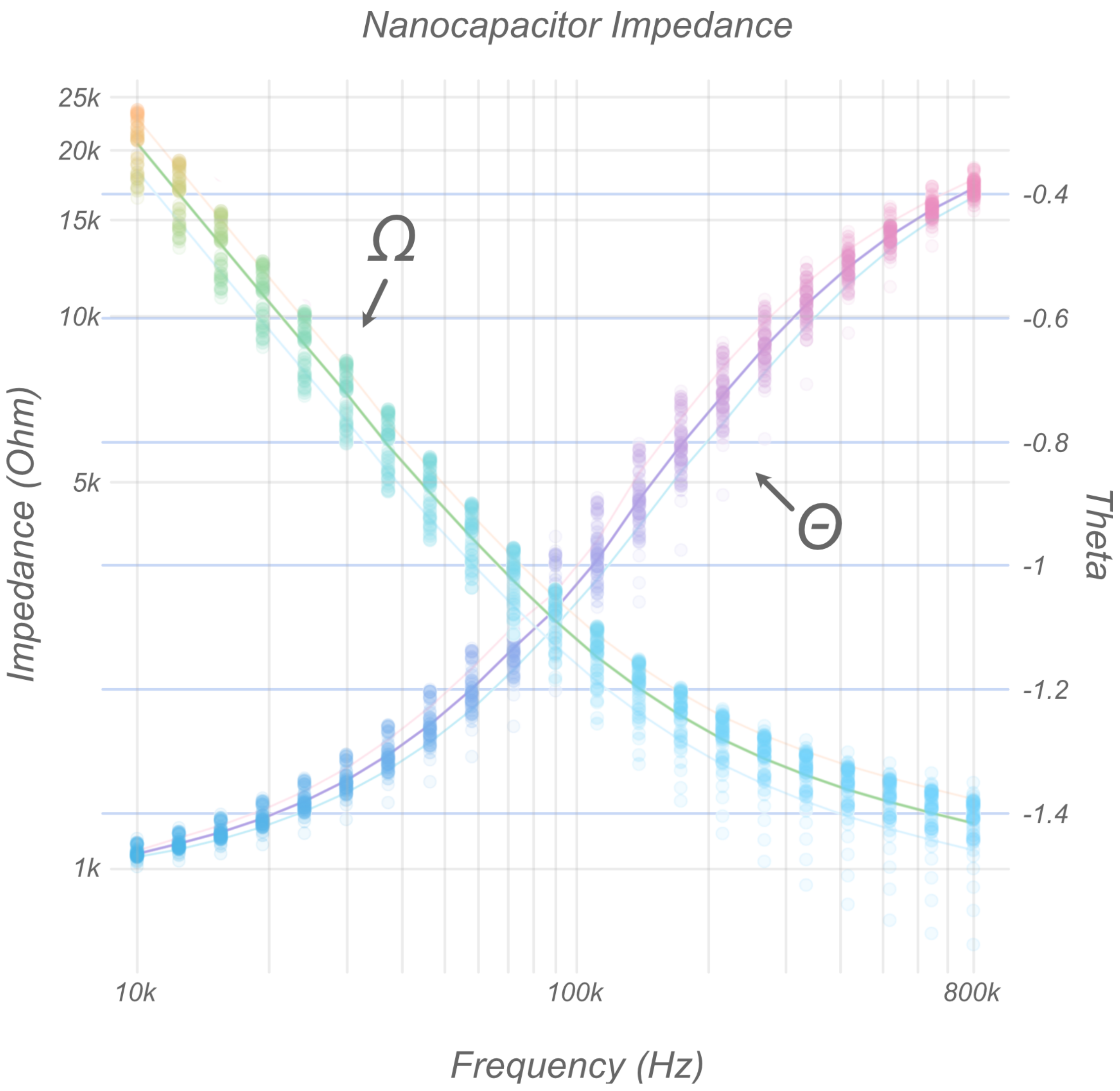


**Supplementary Figure 15: Electrochemical impedance spectroscopy of nanocapacitors.**

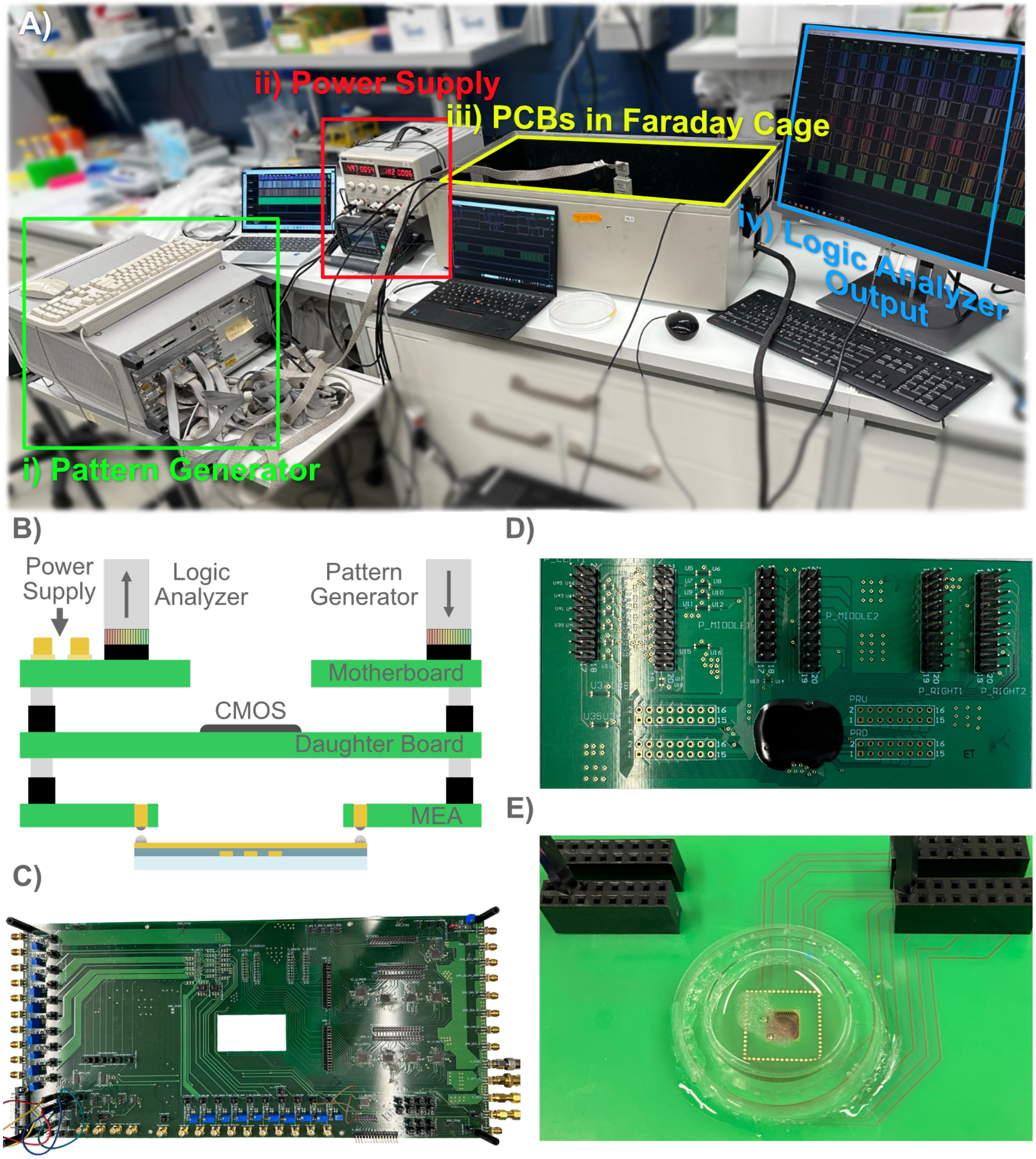


**Supplementary Figure 16: Recording setup. (A)** Layout of a typical recording. An external pattern generator (i) is connected to the motherboard in the faraday cage (iii). External power is supplied (ii) and the data from the experiment is output through logic analyzers to a computer (iv). (B) Cartoon of the PCB stack and the external connections. (C) A picture of the mother board which contains the peripheral connections for running the ASIC. (D) The daughter board which contains the ASIC and connects to the MEA. (E) An assembled MEA with cultured neurons. In contrast to CMOS MEAs, recordings are performed off chip.
